# Epigenetic alterations underlie airway macrophage differentiation and phenotype during lung fibrosis

**DOI:** 10.1101/2020.12.04.410191

**Authors:** Peter McErlean, Christopher G. Bell, Richard J. Hewitt, Zabreen Busharat, Patricia P. Ogger, Poonam Ghai, Gesa Albers, Shaun Kingston, Philip L. Molyneaux, Stephan Beck, Clare M. Lloyd, Toby M. Maher, Adam J Byrne

**Affiliations:** Inflammation, Repair and Development Section, National Heart and Lung Institute, Imperial College, London SW7 2AZ, UK; William Harvey Research Institute, Barts & The London School of Medicine, Queen Mary University of London, Charterhouse Square, London EC1M 6BQ, UK; National Institute for Health Research (NIHR) Respiratory Biomedical Research Unit, Royal Brompton Hospital, Sydney Street, London SW3 6NP, UK; Department of Cancer Biology, UCL Cancer Institute, University College London, Paul O’Gorman Building, 72 Huntley Street, London WC1E 6BT, UK; Hastings Centre for Pulmonary Research and Division of Pulmonary, Critical Care and Sleep Medicine, Keck School of Medicine, University of Southern California, Los Angeles, USA

**Keywords:** Pathogenesis, Monocytes, Epigenetics, DNA methylation, Interstitial lung disease

## Abstract

Airway macrophages (AMs) are key regulators of the lung environment and are implicated in the pathogenesis of idiopathic pulmonary fibrosis (IPF), a fatal respiratory disease with no cure. However, the epigenetics of AMs development and function in IPF are limited. Here, we characterised the DNA-methylation (DNAm) profile of AMs from IPF (n=30) and healthy (n=14) donors. Our analysis revealed epigenetic heterogeneity was a key characteristic of IPF AMs. DNAm ‘clock’ analysis indicated epigenetic alterations in IPF-AMs was not associated with accelerated ageing. In differential DNAm analysis, we identified numerous differentially methylated positions (DMPs, n=11) and regions (DMRs, n=49) between healthy and IPF AMs respectively. DMPs and DMRs encompassed genes involved in lipid (*LPCAT1*) and glucose (*PFKB3*) metabolism and importantly, DNAm status was associated with disease severity in IPF. Collectively, our data identify that profound changes in the epigenome underpin the development and function of AMs in the IPF lung.

## Background

Airway macrophages (AMs) are sentinel innate cells of the lungs contributing to homeostasis and immune response^1^. Ontogeny of AMs is complex encompassing both self-renewing-fetal-derived ‘resident’ and monocyte-derived ‘recruited’ cells^2^. Understanding of AM ontogeny in disease states and aging is contentious and has relied heavily on murine models. However, we recently helped clarify AM ontogeny in humans by identifying that one year post lung transplant, AMs in adults are derived exclusively from recruited peripheral monocytes^3^.

The influence of the local microenvironment in shaping macrophage development and function is increasingly being appreciated^4^. Responses to growth factors or inflammatory mediators can skew macrophage development as exemplified by the pro-inflammatory ‘M1’ and pro-wound healing ‘M2’ paradigm. However, *in vivo* macrophages exhibit tremendous heterogeneity in both health and diseased states^5–7^, indicating a remarkable plasticity.

Key processes in macrophage development are reflected in changes to the epigenome including DNA methylation (DNAm)^8^. Occurring in the context of cytosine-guanine dinucleotides (CpGs), DNAm influences chromatin accessibility, transcription factor (TF) binding and gene expression^9,10^. DNAm represents one of the most stable epigenetic marks and can be measured as a means of assessing the influence of development and diseases on the epigenome. In AMs, DNAm is altered in genetic and environmentally-induced chronic airway diseases^11–13^. Regional differences in lung anatomy also influence DNAm in AMs^14^, suggesting that shaping of AM development in the lung microenvironment comprises an epigenetic component. However, despite recent advances in our understanding of AM ontogeny, the epigenetics of monocyte to macrophage development in the lung and influence of disease on these processes remain limited.

Idiopathic pulmonary fibrosis (IPF) is a deadly respiratory disease of unknown aetiology with heterogeneous cellular and molecular mechanisms^15^. The pathobiology of IPF is characterised by a pro-fibrotic wound-healing cascade that does not resolve, leading to progressive scarring, loss of lung function and ultimately death^16^. Although containing a strong genetic component^17^, the greatest risk factor for IPF is age (median 65 years^18^) and prognosis in IPF is worse than some cancers with a mean survival of 3-5 years^19^. In the IPF lung, AMs exhibit transcriptional^7^, immuno-phenotypic^1^ and metabolic differences^20,21^. Recent studies employing single-cell RNA sequencing (scRNA-Seq) have indicated a transcriptional spectrum of AMs in the IPF lung that reflects facets of both M1 and M2 macrophage paradigm^5–7^. However, despite their emerging role in IPF pathogenesis, the molecular mechanisms underlying transcriptional and other phenotypic characteristics of AMs in IPF are poorly understood.

In the current study we investigated the epigenetics of AMs by undertaking genome-wide DNAm profiling using the Illumina EPIC (850k) arrays. By comparing AMs and other myeloid cell DNA methylomes, we sought to clarify the epigenetics of AM development in the lung. By profiling AMs from healthy and IPF donors, we also sought to determine if changes in the epigenome characterize features of AMs observed in the IPF lung.

## Results

### The DNA methylation profile of AMs is distinct from that of peripheral monocytes or cultured macrophages

Given recent work identifying AMs as monocyte-derived and having characteristics spanning the M1-M2 spectrum of activation, we sought to determine if these changes are also reflected at the epigenetic level by comparing the DNA methylome in AMs and other myeloid cells.

Firstly, we enriched CD206+ AMs, obtained through bronchoalveolar lavage, from healthy (n=14) and IPF (n=30) donors and assayed DNAm using Illumina Methylation EPIC (850k) arrays, which interrogate >850,000 CpGs across the genome with an enrichment for functional loci (promoters and enhancers, Table S1)^22^. These data were then merged with whole genome bisulphite sequencing (WGBS) Blueprint datasets from representative myeloid cell-types including CD14+CD16-‘Classical’ and CD14+CD16+ ‘Other’ monocytes and *in vitro*-derived M0, M1, and M2 macrophages^23^ (Figure 1A). We then identified the top 500 CpGs with a DNAm profile which best discriminated each monocyte and macrophage subtype (see methods) and characterized these as ‘myeloid marker CpGs’ (myld-CpGs, Table S2).

**Figure 1.**
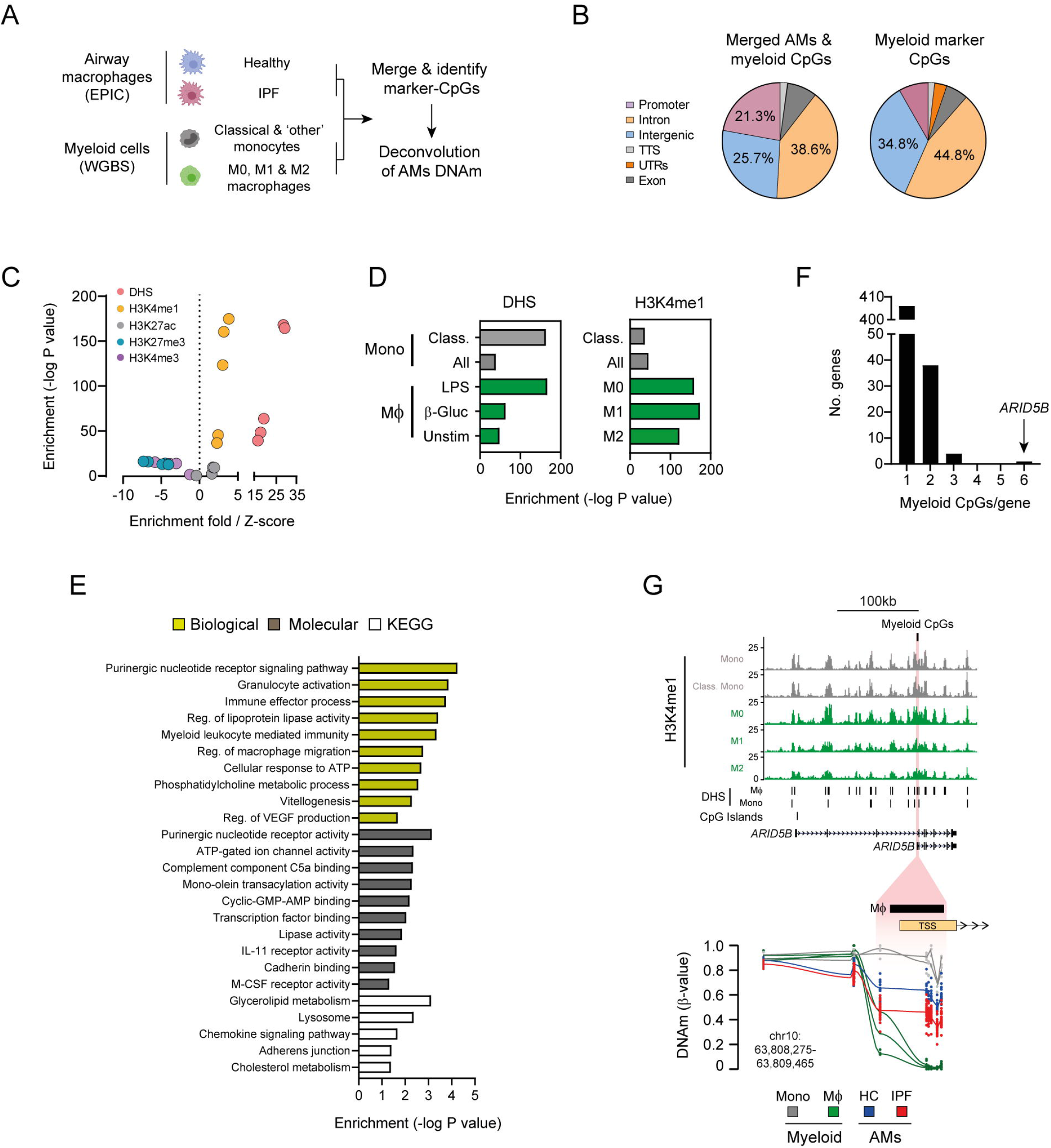
(A) Outline of approach to investigate relationship between airway macrophages (AMs) and other myeloid cell-types (i.e. monocytes and macrophages) DNA methylation (DNAm) as determined by EPIC and whole genome bisulphite sequencing (WGBS) respectively. Cell-type depictions were generated using www.biorender.com. (B) Genomic feature distribution for merged AMs and myeloid cell DNAm datasets and the n=500 myeloid marker CpGs to be used in deconvolution analysis. (C-D) Enrichment of myeloid marker CpGs across histone modifications and DNAse hypersensitivity sites (DHS) identified in myeloid cells. (E) Gene ontology processes and KEGG pathway enrichment analysis for myeloid marker CpGs. (F) Distribution of myeloid marker CpGs per gene. (G) Genome track depicting epigenomic features (H3K4me1 ChIP-Seq, DHS - Blueprint) of myeloid cells across the *ARID5B* loci. The location of n=6 myeloid marker CpGs that cluster at the *ARID5B* variant 2 transcription start site (TSS) is highlighted in red and magnified further below to show DNAm profiles (beta values) across each myeloid cell type and AMs. Lines indicate average DNAm across all of the CpGs assayed in the magnified region.

We found that myld-CpGs reside predominately in intronic and intergenic regions (Figure 1B) that are enriched for other epigenetic features in myeloid cells including histone modifications indicative of poised enhancers (H3K4me1 without H3K27ac) and open chromatin (DNase-I hypersensitivity sites: DHS, Figure 1C-D). Functional enrichment analysis additionally indicated that myld-CpGs encompass a diverse range of receptor signalling, immune cell activation, chemokine and metabolic-related processes and pathways (Figure 1E). Although annotated to n=449 genes, we found AT-Rich Interaction Domain 5B (*ARID5B*), a transcriptional co factor which has been shown to regulate glucose metabolism^24^, contained the most myld-CpGs (Figure 1F, Table S2)

We focused further on *ARID5B* and mining scRNA-Seq datasets from healthy and diseased lung (IPF/COPD) established that *ARID5B* is expressed across immune cells including monocytes and macrophages (Figure S1B). At the *ARID5B* locus we found that the myld-CpGs are clustered at the promoter region of a shorter transcript variant 2 and overlap with DHS and H3K4me1 enrichment (Figure 1G). We then confirmed the expression of the shorter *ARID5B* transcript in AMs (Figure S1C). Finally, closer inspection revealed dramatic changes in DNAm towards the shorter *ARID5B* variant promoter region with AMs exhibiting an intermediate DNAm profile (avg. 50.8%) compared to other myeloid cells (monocytes - avg. 85.4% and macrophages - avg. 0.6%, Table S2). Taken together, these results indicate that the DNAm profile of human AMs is distinct from that of peripheral monocytes/cultured macrophages and we identify *ARID5B* DNAm status as a marker of AM development.

### Changes in the AM-methylome define IPF-AMs

Next, to determine whether the methylome was distinct in each cell-type or disease state, we clustered DNAm profiles for genes with >2 myld-CpGs. Interestingly, our analysis indicated that epigenetic heterogeneity is a feature of IPF, as DNAm profiles across myld-CpGs were distinct when comparing healthy and IPF AMs (Figure S1A). Furthermore, the DNAm of AMs overlapped significantly with other myeloid cells, potentially indicating monocytic origin (Figure S1A).

To clarify this further we performed deconvolution analysis of AM DNAm datasets with mlyd-CpGs (see methods). Deconvolution revealed that while at the epigenetic level healthy and IPF AMs were predicted to be largely a composite of ‘other’ monocytes and M0/M2 macrophages, clustering identified the separation of healthy and IPF AMs (Figure 2A), driven largely by differences in the minor classical monocyte and M1 macrophage fractions (Figure 2B). However, further investigation revealed that the specific differences in subsets was related to donor age (Figures 2C).

**Figure 2:**
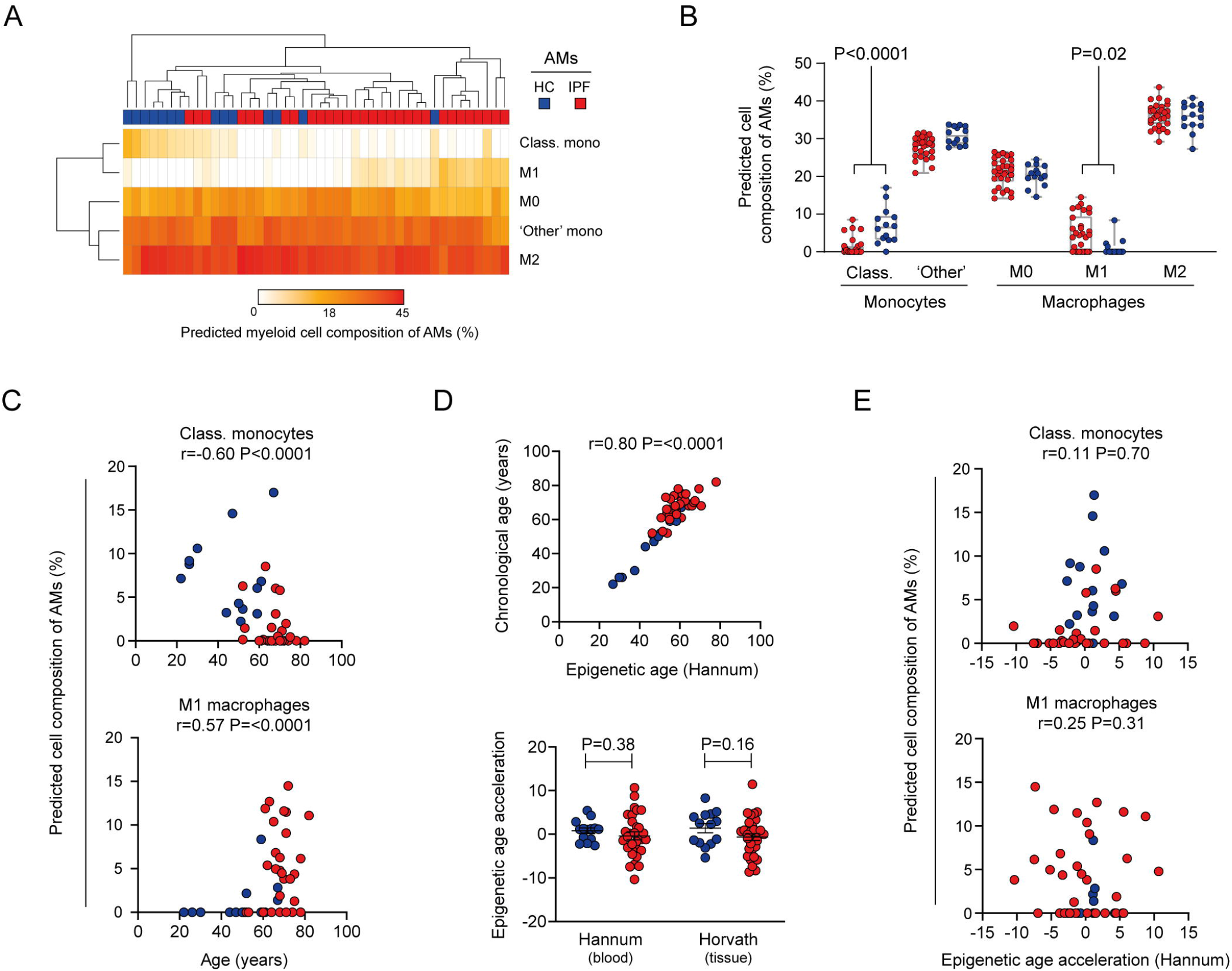
(A) Heatmap depicting predicted myeloid cell composition of airway macrophages (AMs - columns) after deconvolution of DNA methylation (DNAm) profiles with myeloid maker-CpGs generated from reference monocyte and macrophage methylomes (rows). (B) Difference in myeloid cell composition were evident for AMs derived from healthy and IPF donors for ‘classical’ and ‘M1 macrophages’ respectively. P-values determined by one-way ANOVA with Tukey’s correction for multiple testing. (C) Spearman-rank correlation between donor age and composition of AMs DNAm attributed to classical monocytes and M1 macrophages. (D) Epigenetic clock analysis indicating a correlation between chronological and epigenetic age as determined by the Hannum et. al ‘clock’ (top). By comparing residuals from two age-adjusted epigenetic ‘clocks’ (Hannum and Horvath) it was determined IPF AMs exhibited no epigenetic age acceleration compared to AMs from healthy controls. (E) Spearman-rank correlation between epigenetic age acceleration and composition of AMs DNAm attributed to classical monocytes and M1 macrophages.

Because IPF and ageing are linked and many age-related diseases exhibit ‘accelerated’ changes to the epigenome, we next used DNAm ‘clock’ analyses to clarify the contribution of ageing towards the predicted myeloid cell composition of AMs. Epigenetic ‘clocks’ use changes in DNAm to estimate sample donor age and determine if accelerated epigenetic ageing are present (i.e. older age prediction than chronological age) and if disease status is associated with accelerated epigenetic signatures (see Methods). We found that while a strong correlation between chronological and epigenetic age was present, no differences in age-adjusted epigenetic age acceleration was observed between healthy and IPF AMs across either the blood or tissue-derived Hannum and Horvath ‘clocks’ respectively (Figure 2D and S1D). Furthermore, there was no relationship between predicted myeloid cell composition and epigenetic age acceleration (Figure 2E), inferring that myeloid cell composition was a feature of IPF AMs and not the more generalised age-related changes detected by these ‘clocks’. Taken together, these data indicate that epigenetic heterogeneity is present in AMs and is a characteristic of IPF.

### Identification of differentially methylated positions (DMPs) in IPF

We next sought to determine whether AMs DNAm profiles are impacted during IPF and to identify the impact of myeloid cell composition in these analyses. Initial principal component analysis indicated a separation of donors by disease group (Figure S2A) with myeloid cell composition and donor age being comparable drivers of variance within the dataset (Figure S2D). We then undertook analysis to identify DMPs in IPF and observed a dramatic impact when adjusting for myeloid cell composition in addition to other study covariates (Figure 3A and methods). This was equally evident when investigating direction of DNAm change in IPF with all myeloid-adjusted DMPs identified (n= 11) losing DNAm compared to healthy controls (Figures 3B, S2C-D and Table S3).

**Figure 3:**
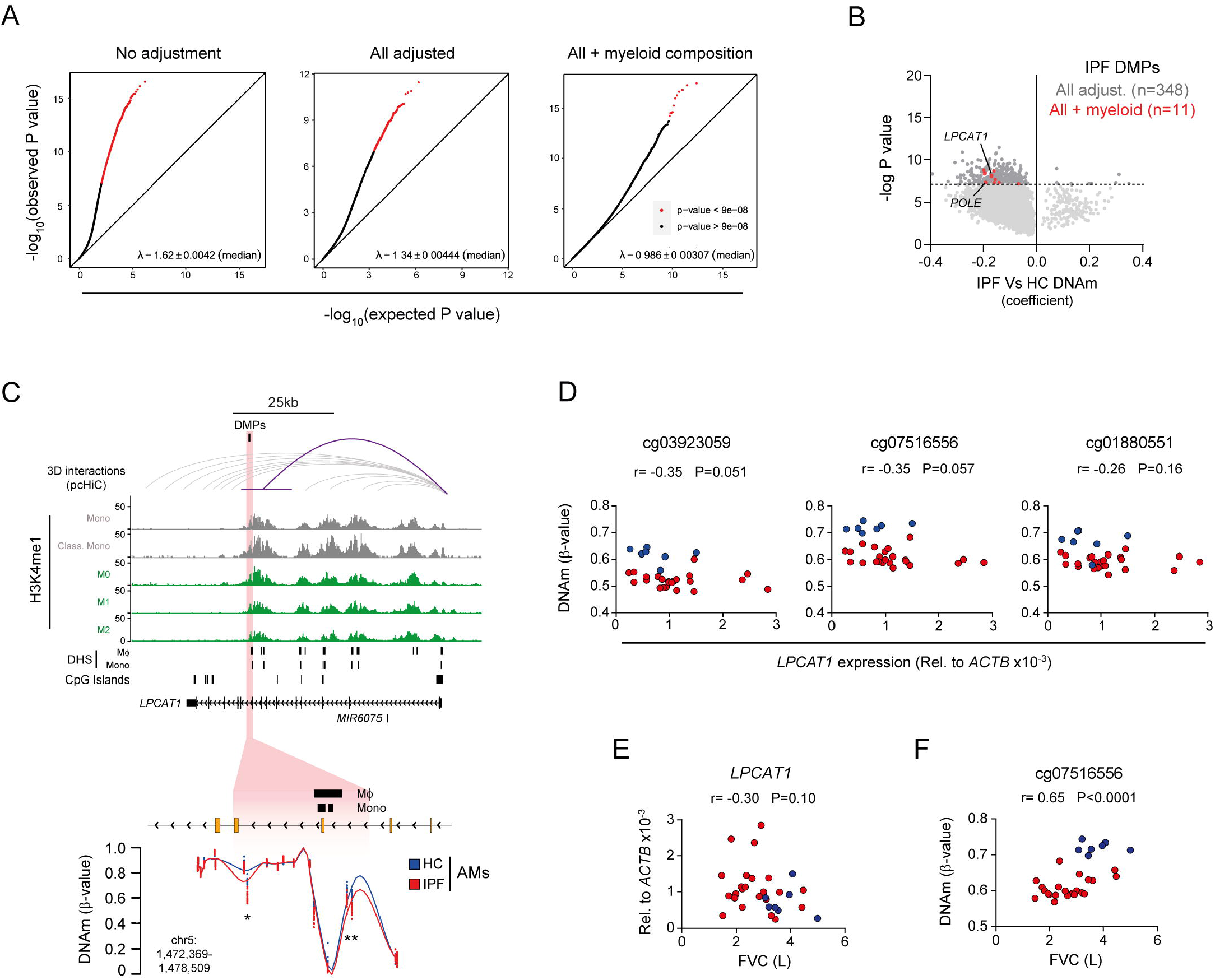
(A) Quantile-quantile plots depicting the impact of adjustment for myeloid cell composition in addition to other study covariates on identification of differentially methylated positions (DMPs) in IPF. Those DMPs reaching the epigenome-wide significance (EWAS) threshold of P<9×10^-8^ are highlighted in red. (B) Volcano plot depicting impact of myeloid cell-adjustment and direction of DNA methylation (DNAm) changes of DMPs in IPF. Dashed line represents EWAS P-value threshold. (C) Genome track depicting epigenomic features (H3K4me1 ChIP-Seq, DNase-I hypersensitivity - DHS - Blueprint) of myeloid cells across the *LPCAT1* loci. Regions interacting with the *LPCAT1* promoter in 3D as determined by promoter-capture HiC (pcHiC) are indicated in grey. Interactions with frequency threshold >5 in myeloid cells are highlighted in purple. The location of n=3 intronic IPF DMPs are highlighted in red and magnified further below to show methylation profiles (beta values) of healthy and IPF AMs across the respective CpGs (*). Lines indicate average methylation across all of the CpGs assayed in the magnified region. (D) Relationship between DNAm of IPF DMPs and gene expression for *LPCAT1* across Healthy (blue) and IPF (red) donor AMs. (E-F) Relationship between gene expression (E), methylation of an IPF-associated DMP (F) for *LPCAT1* and forced vital capacity (FVC).

IPF DMPs were either intronic (n=9) or intergenic (n=2) and occurred in regions enriched for open chromatin in myeloid cells (DHS, Figure S2E). We found n=3 IPF DMPs clustered at Lysophosphatidylcholine Acyltransferase-1 (*LPACT1*), an enzyme which mediates the conversion of lysophosphatidylcholine to phosphatidylcholine^25^ (Figure 3B). Mining of scRNA-Seq data indicates that *LPCAT1* is expressed across monocytes and macrophages in the lung and we confirmed these findings in our study AMs (Figure S1B-C).

Although *LPCAT1* DMPs are intronic, distal regions can influence gene expression through 3D interactions. To investigate this further, we used promoter capture HiC (pcHiC) data to investigate the relationship between IPF DMPs and 3D interactions in myeloid cells^26^. Remarkably, while the *LPCAT1* promoter interacted with other genes/regions specifically in monocytes (Figure S2F), the strongest interaction occurred with the region containing the IPF DMPs (Figure 3C). We identified a correlation between IPF DMPs methylation and gene expression occurred only for *LPCAT1* (Figure 3D) and not any other interacting genes (Figure S2G).

Finally, we investigated the relationship between *LPCAT1* and clinical features of IPF (Table S1) and found that while no relationship was present for gene expression (Figure 3E), there was a strong correlation between methylation and forced vital capacity (FVC), a measure of disease severity and progression in IPF^27^ was evident (Figure 3F). Taken together these data suggest AMs similarity to monocytes on the epigenetic and higher order 3D-interaction level and a function of DNAm of AMs in IPF pathogenesis.

### Identification of differentially methylated regions (DMRs) in IPF

Given the clustering of IPF DMPs, we next conducted analysis to identify DMRs in IPF. Similar to DMP analysis, we saw a reduction in total DMRs identified after adjusting for myeloid cell composition (Figure 4A). However, we found n=49 myeloid-adjusted DMRs which included regions both gaining and losing DNAm compared to healthy controls (Table S4). We also found n=2 DMRs which encompassed the previously identified DMPs of *LPCAT1* and DNA Polymerase Epsilon, Catalytic Subunit (*POLE*, Figure 3B).

**Figure 4:**
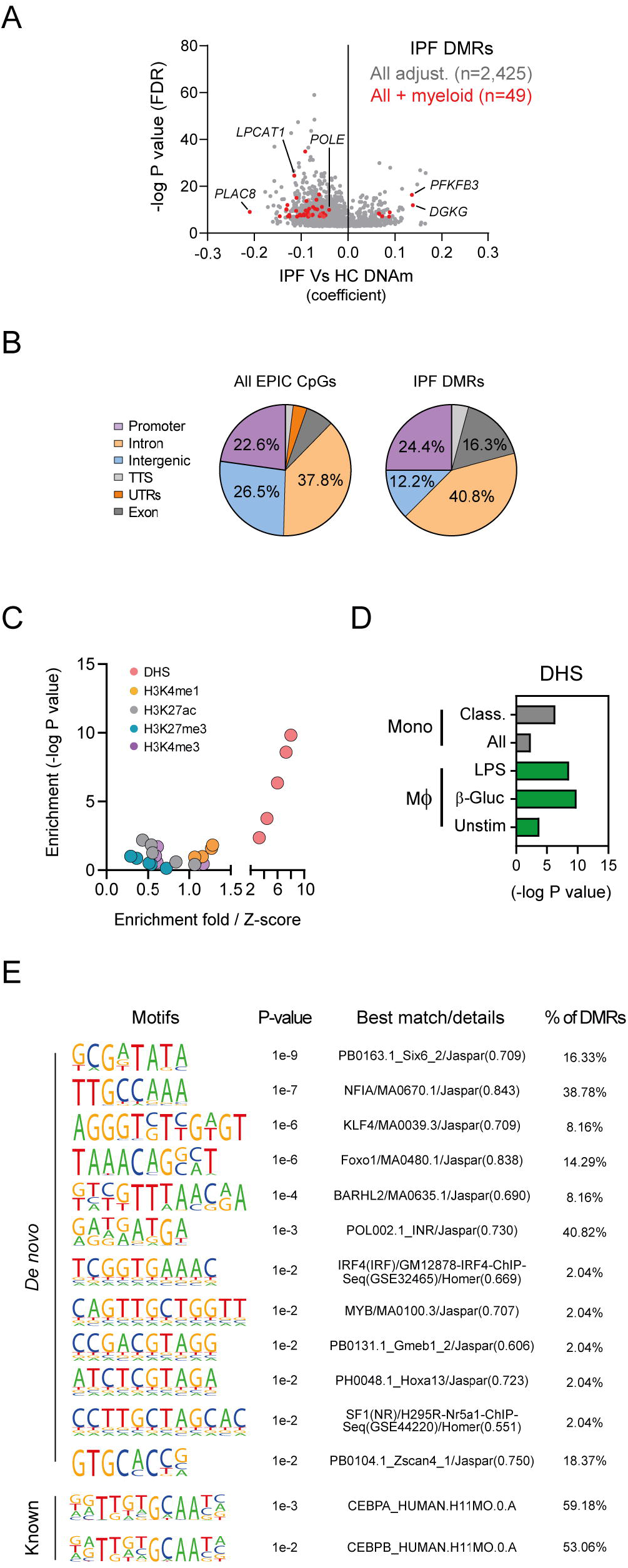
(A) Volcano plot depicting impact of myeloid cell-adjustment and direction of DNA methylation (DNAm) changes of differentially methylated regions (DMRs) in IPF. (B) Genomic feature distribution for all EPIC array CpGs and those encompassed by IPF DMRs. (C-D) Enrichment of IPF DMRs across histone modifications and DNAse-I hypersensitivity sites (DHS) in monocytes and macrophages. (E) DNA motif enrichment in IPF DMRs.

IPF DMRs were distributed across various genomic features including promoters, introns and exons (Figure 4B), occurred in regions enriched for open chromatin in myeloid cells (Figures 4C-D) and were more likely linked in 3D to distal genes and regions (Figure S3A-C). We additionally found motifs matching TF’s previously implicated in macrophage polarisation to be enriched in IPF DMRs (e.g. KLF4, FOXO1, Figure 4E) and the subsequent cell-type expression profiles of TF encoding genes across lung immune cells (Figure S3D).

We then undertook functional enrichment analysis and found enrichment across various processes and pathways pertinent to macrophage biology (e.g. extravasations) and IPF pathogenesis (e.g. platelet activation, response to wound healing; Figure S4A).

To gain a better insight into the biological implications of changes in DNAm, we undertook additional protein-protein interaction analysis and found that DMR-associated genes form central hubs in a large interconnect network (Figure 5A and Figure S4B). We refined our analysis further and undertook functional enrichment of networks by DNAm status and found hub genes gaining DNAm in IPF predominately encompass metabolic processes whilst those losing DNAm play a role in processes and pathways pertinent to macrophage biology and fibrogenesis (e.g. phagocytosis, cell proliferation and TGF-β signalling, Figure 5B).

**Figure 5:**
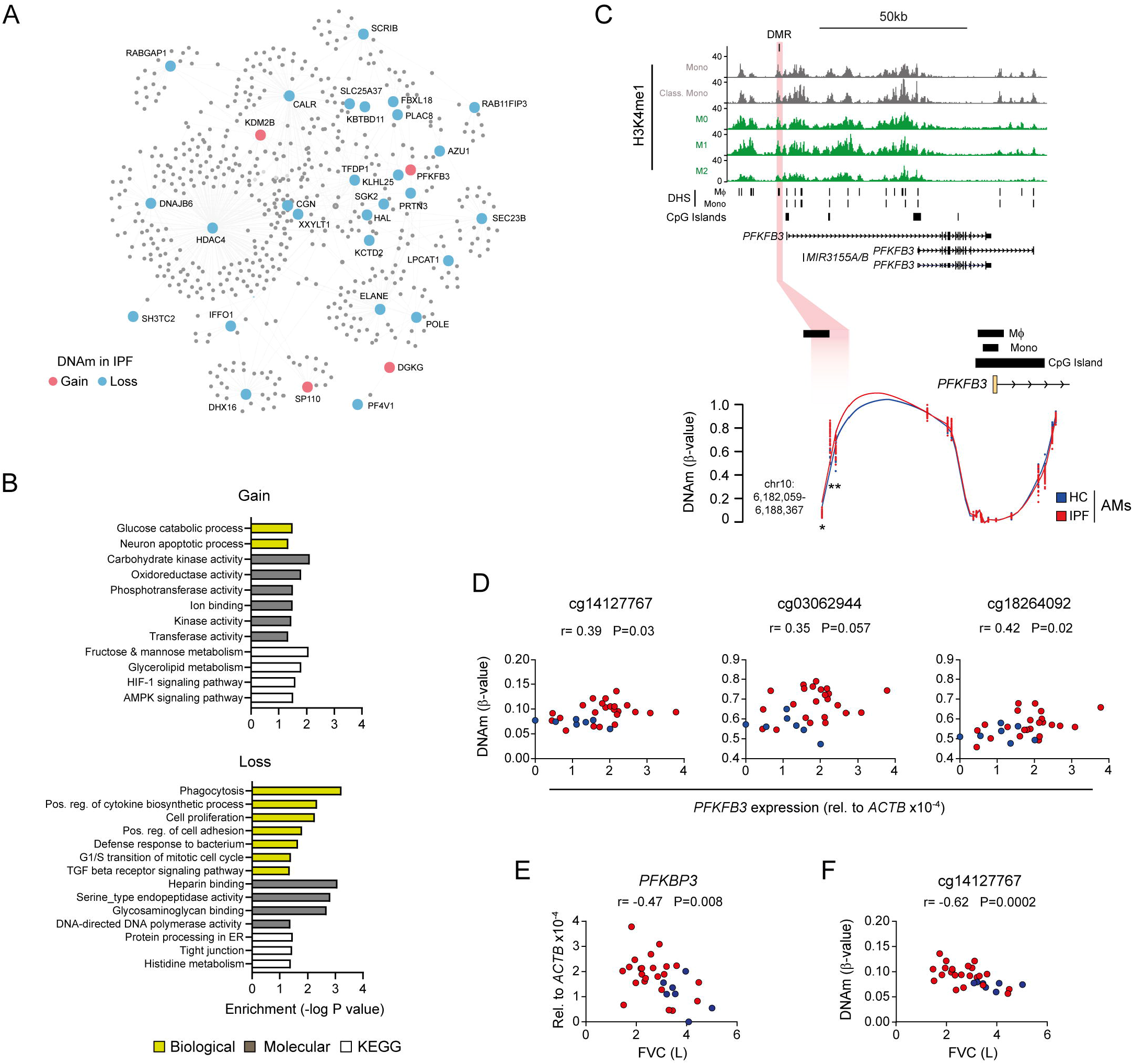
(A) Network depicting protein-protein interactions of DMRs-associated genes. (B) Gene ontology processes and KEGG pathway enrichment analysis for DMR-associated genes gaining or losing DNA methylation (DNAm) in IPF. (C) Genome track depicting epigenomic features (H3K4me1 ChIP-Seq, DNase hypersensitivity - DHS - Blueprint) of myeloid cells across the *PFKFB3* loci. The location of the IPF DMR is highlighted in red and magnified further below to show methylation profiles (beta values) at the associated n=3 CpGs (*) across healthy and IPF AMs. Lines indicate average methylation across all of the CpGs assayed in the magnified region. (D) Relationship between DNAm of IPF DMPs and gene expression for *PFKFB3* across Healthy (blue) and IPF (red) donor AMs. (E-F) Relationship between gene expression (E), methylation of DMR-associated CpG (F) for *PFKFB3* and forced vital capacity (FVC).

Given that work from our lab has identified an altered state of AM metabolism in IPF^20,21^, we focused on 6-Phosphofructo-2-Kinase/Fructose-2,6-Biphosphatase 3 (*PFKB3*), a potent driver of glycolysis^28^ and found the IPF related DMR overlapped H3K4me1 and DHS enrichment and was located in an intergenic region, upstream of the *PFKB3* promoter. Remarkably, all AMs exhibited a complete loss of methylation for CpGs at the *PFKB3* TSS (Figure 5C). We found *PFKB3* exhibited differential expression between healthy and IPF AMs (Figure S1C) and subsequently confirmed a correlation between DNAm and gene expression for two of the 3 CpGs encompassing the *PFKB3* DMR (Figure 5D). We additionally identified relationships between *PFKB3* gene expression and methylation with severity of IPF as determined by FVC (Figure 5E-F). Taken together these data strongly suggest that changes in the epigenome underpin the distinct metabolic phenotype observed in AMs isolated from IPF lung and their contribution towards disease pathogenesis.

## Discussion

AMs are key regulators of the lung environment and are implicated in the pathogenesis of lung fibrosis. By comparison to reference myeloid cells, we determined that epigenetic heterogeneity is present in AMs and, furthermore, is a characteristic of IPF (Figure 2). While identified computationally, our findings mirror scRNA-Seq studies of the IPF lung where AMs exhibit transcriptional heterogeneity^5–7^. Differences in myeloid cell composition also suggests that the IPF lung influences monocyte to macrophage developmental trajectories. Interestingly, transcriptomic signatures reflective of blood monocytes are already altered in association with IPF severity^29–31^, potentially indicating that the effects of IPF extends across tissue compartments rather than being isolated to the lung. As such, blood monocytes from individuals with IPF may already be ‘primed’ towards a particular macrophage lineage (e.g. M1-like). With advancements in single-cell epigenomics, future studies of IPF should seek to generate matched transcriptome and epigenomic datasets across blood and lung to comprehensively address the molecular events and influence of IPF on AM developmental trajectories.

In the absence of single-cell data, computational deconvolution of epigenetic data is essential to decipher disease effects within samples consisting of mixed cell populations^32^. Even though we had enriched AMs based on cell surface expression of CD206, we identified epigenetic heterogeneity (Figure 2A) and found a tremendous impact of myeloid cell composition in identification of DMPs and DMRs in IPF (Figure 3B and Figure 4A). Our work has implications for previous studies of DNAm in IPF that have largely assayed whole lung tissue in ‘bulk’ without accounting for cell-type heterogeneity^33,34^. Furthermore, our study employed a genome-wide approach, providing better insights into the influence of IPF on the wider epigenome than previously conducted gene-specific studies in this disease area^35^.

While the identification of epigenetic heterogeneity in AMs was important for deciphering DNAm changes in IPF, we additionally found that donor age correlated with this predicted heterogeneity (Figure 2C). To address the potential interaction of heterogeneity and age, we conducted epigenetic ‘clock’ analysis as these signatures are actively being explored for possible novel age-related disease insights^36^. However, we found no differences in age acceleration between healthy and IPF AMs or relationship to predicted myeloid cell composition (Figure 2D-E). These findings are in contrast to many other age-related diseases^37^ and indicate that whilst IPF predominately occurs in later decades, the DNAm changes detected are likely to be specific to IPF rather than representing epigenome changes occurring as a consequence of otherwise ‘healthy ageing’^38^. Although these analyses and other adjustments for donor age in differential analysis revealed influence of IPF on AMs DNAm, IPF and age remain inexplicably linked. Future studies of epigenetics in IPF should therefore strive to include age and sex matched healthy controls.

By attempting to clarify the epigenetic events related to macrophage development in the lung, we found that DNAm patterns which discriminate myeloid cell-type occur largely in intronic and intergenic regions (Figure 1B). This supports previous work indicating epigenomic changes during immune cell lineage commitment occurs within non-coding regions^39^. However, we identified DNAm intragenically within *ARID5B* at the promoter locus for a shorter transcript variant 2 as a mark of monocyte to macrophage development (Figure 2G). *ARID5B* is a chromatin modifier that acts as a transcriptional coactivator by removing repressive histone modifications^40^. Additionally, *ARID5B* has been linked to adipogenesis^41^ and metabolism in hepatocytes^42^ and natural killers cells, where altered DNAm particularly of the short transcript variant 2 characterized a HMCV+ adaptive NK cell substype^43^. More relevant to this study was work using a multi-omics approach that identified *ARID5B’*s association with atherosclerosis in CD14+ blood monocytes and implicated 3D interactions in linking intronic *ARID5B* DNAm (and other regulatory regions) with the *ARID5B* promoter^44^.

We identified n=11 EWAS DMPs in IPF and found most changes were clustered within an intronic region of *LPCAT1* (Figure 3C). *LPCAT1* is an evolutionarily conserved enzyme that is involved in phospholipid metabolism and performs a key role in surfactant production in alveolar type 2 cells^45^ and inflation of lungs upon birth^46^. Recent work has implicated *LPCAT1* with aberrant metabolism and plasma membrane remodelling in cancer, helping to establish functional links between genetic alterations and tumour growth^47^. In IPF, reduction of *LPCAT1* gene expression was shown to characterise subsets of IPF-specific airway epithelial cells^48^.

Similar to work in blood monocytes^44^, we investigated whether integrating 3D interactions could help elucidate the potential impact of changes in DNAm on gene expression^26^. Remarkably, we found the *LPCAT1* promoter ‘self-interacted’ with regions containing IPF DMPs (Figure 3C) and over half of all DMRs were linked in 3D to other genomic regions in myeloid cells (Figure S3C). While these data suggest that similar to DNAm, AMs share a higher order chromatin structure similar to monocytes and macrophages, bias from the EPIC array design needs to be taken into consideration. Future studies should therefore aim to determine 3D interactions in AMs from healthy and IPF donors empirically.

Work from our lab has indicated the crucial role of immunometabolism of AMs in IPF^20,21^. We identified IPF DMRs encompassed genes and networks of lipid, iron and glycolytic metabolic processes (Figure 5B and S4A-B). Indicative of the impact of changes in DNAm was the DMR located at *PFKFB3* (Figure 3C), an enzyme responsible for the synthesis and degradation of fructose 2,6-bisphosphate, a key regulator of glycolysis. In macrophages, work has identified an important role of *PFKFB3* with plasticity and the M1-phenotype in liver fibrosis^49^ and with HIF-1α in driving glycolytic flux and maintaining cell viability under hypoxic and inflammatory conditions^50^. Given the composition of AMs in IPF was more ‘M1-like’ (Figure 2B) and the progressive remodelling of the IPF lung results in an inflammatory hypoxic environment, these results raise the question of whether development and epigenetic changes identified in IPF AMs are a cause or consequence of the fibrotic milieu of the IPF lung.

In conclusion, our study has identified a role of aberrant epigenetic regulation of AMs, independent of ageing alone, which appears to be involved in IPF pathogenesis. Our study provides a foundation for further investigations to clarify the role of epigenetics during monocyte to macrophage development in the healthy and diseased airways. Furthermore, our data highlight the possibility that therapeutic agents targeting epigenetic modification may have a role in the treatment of IPF.

## Methods

### Patient recruitment and sample collection

Study donors underwent bronchoscopy and collection of BAL as outlined previously^20,21^. All study donors provided written informed consent to participate in the study, which was approved by the research ethics committee (10/HO720/12, 15/LO1399 and 15/SC/0101). Clinical characteristics of donors are outlined in Table S1. Differences in donor data were determined by Mann-Whitey or Chi-square Test for quantitative and categorical data respectively using GraphPad Prism v.8.4.2.

### AMs enrichment

CD206+ AMs were enriched from donor BAL using the magnetic-based MACS^®^ system (Miltenyi Biotech, Germany) and fluorescent activated cells sorting (FACS) as outlined previously^3,20,21^. Briefly, for MACS-based enrichment, BAL cells (1 × 10^7^) were incubated with human Fc-block (BD Biosciences, USA) and human CD206 APC-Cy7 (clone 15-2, BioLegend, USA). CD206+ cells were enriched in MACS LS magnetic separation column and MidiMACS™ magnet. Cell counts were determined using a haemocytometer and trypan blue live/dead exclusion.

For FACS-based enrichment BAL cells were washed and incubated with near-infrared fixable live/dead stain (Life Technologies Inc.) as per the manufacturer’s instructions. After incubation with human Fc block (BD Pharmingen, Inc.), surface staining was performed with the following antibodies (fluorophore followed by clone in parentheses); CD45 (PE-Texas Red, H130), CD3 (FITC, OKT), TCR-β (BV421, IP26), CD206 (PercpCy5.5, 15.2). Cell sorting was carried out on Aria III (BD Biosciences) and AMs defined as live, CD45^+^CD3^-^TCR^-^CD206^+^ cells.

### DNA/RNA extraction and EPIC methylation arrays

Nucleic acids were extracted from cells using the AllPrep Mini Kit (QIAGEN, Germany). DNA quality and quantity were assessed using Genomic DNA ScreenTape and TapeStation System (Agilent, USA). DNA was submitted to the UCL Genomics Core facility where bisulphite conversion, hybridization and scanning of Infinium MethylationEPIC BeadChip Arrays (Illumina, USA) were performed according to Illumina recommendations.

### Array QC and pre-processing

We employed RnBeads 2.0 pipeline^51^ for methylation array preprocessing. Briefly, quality control (QC) metrics were generated and samples passing QC (e.g. bisulphite conversion efficiency) were pre-processed to remove probes with a detection P value <0.01, directly overlapping SNPs, those with SNPs within 3nt of the 3’end, cross-reactive with multiple locations and those located on sex chromosomes^52^. Processed data was then normalized using the *Dasen* function implemented from the wateRmelon package^53^. Following QC and pre-processing, 784,669 probes for each n=44 samples remained for downstream analysis.

### Myeloid marker CpGs and deconvolution analysis

Whole genome bisulphite sequencing (WGBS) data for Blueprint methylomes (2016 release)^23^ were accessed through the RnBeads methylome resource (https://rnbeads.org/methylomes.html). Samples representing the myeloid cell compartment (venous blood monocytes and macrophages) and derived from donors of age comparable to our study population (i.e. >50 years) were selected (Table S5) and WGBS data pre-processed through the RnBeads pipeline as outlined above.

EPIC array and WGBS data were then merged (custom scripts available upon request) resulting in 298,945 CpGs across each n=44 CD206+ and n=13 reference methylomes. The top 500 most variable CpGs were then used to identify ‘myeloid-marker CpGs’ and subsequently deconvolute and predict myeloid cell composition of AMs using the Houseman method^54^ implemented in RnBeads. Heatmaps were produced using Morpheus (https://software.broadinstitute.org/morpheus).

### Differential methylation

DMP analysis was performed using the meffil R pipeline^55^ which implements standard and rigorous Illumina DNA methylation array QC, normalisation and subsequently Epigenome-wide Association Study (EWAS) analysis. In addition to preprocessing outline above, standard meffil QC parameters were employed to detect poor probes and/or samples for removal. Zero outliers were detected based on deviations from mean values for control probes and 6 Principal Components (PCs) were assessed to be needed to adjust for technical effects.

In total four IPF Vs Healthy EWAS were run: (i) No adjustment for covariates (ii) adjustments for ‘All’ covariates: age, sex, smoking history, FACS/MACS enrichment; (iii) adjustments for ‘All + myeloid composition’: age, sex, smoking history, FACS/MACS enrichment with addition of predicted monocyte and M0, M1, M2 macrophage composition and (iii) - as outlined for (ii) but Phenotype randomised retaining all other covariate information consistent (data not shown). Quantile-quantile (QQ) plots were generated and inspected for EWAS QC and p-value inflation. A robust genome-wide significance threshold of P value < 9×10^-8^ as detailed by Mansell et al.^56^ was employed to identify significant DMPs. DMRs were identified using DMRcate^57^ after covariate adjustments outlined above. DMRs contained >3 CpGs and were ranked based on min-smoothed false discovery rate (FDR P<0.05).

### Genomic and epigenomic feature enrichment

Genomic feature distribution and annotation of myeloid marker CpGs, DMPs, DMRs and all EPIC probes were identified using HOMER^58^ (*annotatePeaks.pl*). Enrichment across epigenomic features from monocytes and macrophages derived from Blueprint consortia were conducted with eForge 2.0^59^ and EpiAnnotator^60^. For Epiannotator analysis, genomic coordinates were firstly converted from hg19 to hg38 using the *CrossMap BED* tool implemented in Galaxy Europe sever^61^.

H3K4me1 ChIP-Seq data was accessed through the International Human Epigenomics Consortium portal (https://epigenomesportal.ca/ihec/grid.html). All Blueprint H3K4me1 datatracks for ‘monocytes’, ‘CD14-positive, CD16-negative classical monocyte’, ‘macrophage’, ‘inflammatory macrophages’ and ‘alternatively activated macrophage’ were exported and merged in the UCSC genome browser. Additional DNase-seq data was obtained for ‘monocytes’ and ‘macrophages’ using ChIP-Atlas^62^.

### 3D interactions

Promoter capture Hi-C (pcHiC)^26^ was used to investigate relationships between differential DNAm and 3D architecture. pcHi-C data was accessed (https://osf.io/u8tzp/files/) and overlapped with DMPs and DMRs using bedtools. To address the potential biasing of promoter-based capture and prominence of CpGs/EPIC probes at gene promoters, we selected only ‘other-end’ (OE’s) interactions overlapping DMPs/DMRs. Unique ‘baits’ of overlapping OE’s with interactions >5 in monocytes and macrophages where subsequently used to identify genes linked in 3D to differential DNAm. Enrichment of overlap was determined using Chi-square test with Yates correction using GraphPad Prism v.8.4.2.

### Functional enrichment

Gene ontology processes and KEGG pathway enrichment of DMR-associated genes was conducted using *goregion* function implemented in DMRcate. Additional protein-protein interaction networks and functional enrichment were identified using NetworkAnalyst 3.0^63^ and the IMEx Interactome database.

### Epigenetic clock analysis

The minfi R package^64^ was used to extract array data and the recommended probe-type normalization for clock analysis was performed (preprocess = Noob). A subset of the 30,084 CpGs were extracted from the total array probe set using the datMiniAnnotation3.csv file for Advanced Analysis utilising the DNAm age calculator (https://dnamage.genetics.ucla.edu)^65^. Due to differences in the 850k array, 2,552 CpGs are not included in this list. A sample annotation file including donor chronological age, sex, and tissue type was also included. Age-adjusted epigenetic age acceleration across the major blood cell-type-derived Hannum^66^ and pan tissue-Horvath^65^ clocks was calculated for each donor (n=44 total) and subsequently compared between IPF and healthy donors and myeloid cell composition. P values were determined via Mann-Whitney Test using GraphPad Prism v.8.4.2.

### Motif Enrichment

HOMER was used to conduct motif enrichment in DMRs (*findMotifsGenome.pl-size given*). For known motifs the HOCOMOCO v11 database was used^67^

### Diseased lung scRNA-Seq data mining

We accessed lung immune cell scRNA-Seq data via the IPF Cell Atlas (www.ipfcellatlas.com) and utilized the Kaminski/Rosas dataset which includes samples from healthy controls and donors with IPF and COPD^5^.

### Quantitative real-time PCR (qPCR)

Gene expression was performed as outlined previously^20,21^. Taqman probes used in this study were purchased from Thermo scientific: *LPCAT1* (Hs00227357_m1), *SLC12A7* (Hs00986431_m1), *SLC6A3* (Hs00997374_m1), *ARID5B* (Hs01382781_m1), *PFKFB3* (Hs00998698_m1). Spearman rank correlations between differential DNAm, clinical variables and expression were identified using GraphPad Prism v.8.4.2.

### Data availability

All EPIC methylation array data has been deposited on Gene Expression Omnibus (GSE159655).

## Acknowledgments

The authors would like to acknowledge the support of the Imperial College South Kensington flow cytometry facility, particularly Ms. Jane Srivastava and Dr. Jessica Rowley. We wish to thank Dr Mark Kristiansen and Gaganjit Kaur Madhan at University College London Genomics for processing EPIC arrays. We thank Dr Michael Scherer at Department of Genetics/Epigenetics, Saarland University, Germany for advice with RnBeads analysis and providing additional code.

## Abbreviations

AMs: Airway macrophages
CpG: Cytosine-guanine dinucleotides
FVC: Forced vital capacity
HC: Healthy control
DHS: DNase-I hypersensitivity sites
DNAm: DNA methylation
DMPs: Differentially methylated positions
DMRs: Differentially methylated regions
IPF: Idiopathic pulmonary fibrosis
Myld-CpGs: Myeloid marker CpG dinucleotides
pcHiC: Promoter capture HiC
scRNA-Seq: Single-cell RNA sequencing
WGBS: Whole genome bisulphite sequencing

---

**Figure S1** (A) Heatmap depicting DNA methylation (DNAm) profiles across myeloid cells (i.e. monocytes and M0, M1, M2 macrophages) and AMs from healthy and IPF donors for genes containing >2 myeloid marker CpGs (Table S2).

(B) Disease origin and cell-type expression of genes highlighted in this study in single cell RNA-Seq data from the IPF lung. Gene expression is projected over UMAP representation of all disease/cell-types with brighter colours indicating more expression. Full data available at www.ipfcellatlas.com.

(C) Gene expression of genes highlight in this study in AMs from healthy and IPF donors as determined by qPCR. * P<0.05, ** P<0.05 Mann-Whitey Test.

(D) Spearman-rank correlations for chronological and epigenetic age (right) and for epigenetic age acceleration and composition of AMs DNAm attributed to classical monocytes and M1 macrophages (left) as determined by the Horvath et. al ‘clock’.

**Figure S2** (A) PCA plot of normalized DNA methylation (DNAm) data for all CpGs prior to differential analysis indicating separation of healthy and IPF AMs.

(B) Sources of variation in DNAm dataset across the first 5 principal components for all CpGs. Associated P values were generated through RnBeads pipline.

(C-D) Plots depicting changes in DNAm (beta value) for healthy and IPF donors across Intronic (C) and intergenic (D) DMPs (Table S3).

(E) Enrichment of IPF differentially methylated positions (DMPs) across DNAse hypersensitivity sites (DHS) in monocytes and macrophages.

(F) All regions interacting with the *LPCAT1* promoter in 3D as determined by promoter-capture HiC (pcHiC). In addition to self-interacting with the region containing IPF DMPs (purple, Figure 2E), *LPCAT1* additionally interacts with the promoters of *SLC6A3* and *SLC12A7*. Dashed line indicates interactions with frequency threshold >5.

(G) Relationship between DNAm of IPF DMPs and gene expression for *SLC6A3* across Healthy (blue) and IPF (red) donor AMs. No *SLC12A7* gene expression was detected in AMs.

**Figure S3:** (A) Proportion of unique promoter-capture HiC (pcHiC) ‘other ends’ containing any EPIC CpGs or those identified as differentially methylated positions (DMPs) or DMRs in IPF.

(B) Proportion of all EPIC, DMPs and DMRs that are linked in 3D as determined by pcHiC. **** P<0.0001, Chi-square test with Yates correction versus all interactions containing EPIC CpGs.

(C) Number of 3D interactions per IPF DMRs (Table S4).

(D) Cell-type expression of transcription factor genes with enriched motifs in IPF DMRs in single cell RNA-Seq data from the IPF lung. Gene expression is projected over UMAP representation of all cell-types with brighter colours indicating more expression. Full data available at www.ipfcellatlas.com.

**Figure S4:** (A-B) Gene ontology processes and KEGG pathway enrichment analysis for all DMR-associated genes (A) and all those forming protein-protein interactions regardless of DNAm status in IPF (B).

## References

1 Byrne, A. J., Maher, T. M. & Lloyd, C. M. Pulmonary Macrophages: A New Therapeutic Pathway in Fibrosing Lung Disease? Trends in Molecular Medicine 22, 303–316, doi:https://doi.org/10.1016/j.molmed.2016.02.004 (2016).

2 Misharin, A. V. et al. Monocyte-derived alveolar macrophages drive lung fibrosis and persist in the lung over the life span. J Exp Med 214, 2387–2404, doi:10.1084/jem.20162152 (2017).

3 Byrne, A. J. et al. Dynamics of human monocytes and airway macrophages during healthy aging and after transplant. The Journal of Experimental Medicine 217, doi:10.1084/jem.20191236 (2020).

4 Lavin, Y. et al. Tissue-Resident Macrophage Enhancer Landscapes Are Shaped by the Local Microenvironment. Cell 159, 1312–1326, doi:https://doi.org/10.1016/j.cell.2014.11.018 (2014).

5 Adams, T. S. et al. Single-cell RNA-seq reveals ectopic and aberrant lung-resident cell populations in idiopathic pulmonary fibrosis. Science Advances 6, eaba1983, doi:10.1126/sciadv.aba1983 (2020).

6 Habermann, A. C. et al. Single-cell RNA sequencing reveals profibrotic roles of distinct epithelial and mesenchymal lineages in pulmonary fibrosis. Science Advances 6, eaba1972, doi:10.1126/sciadv.aba1972 (2020).

7 Reyfman, P. A. et al. Single-Cell Transcriptomic Analysis of Human Lung Provides Insights into the Pathobiology of Pulmonary Fibrosis. American Journal of Respiratory and Critical Care Medicine 199, 1517–1536, doi:10.1164/rccm.201712-2410OC (2018).

8 Wallner, S. et al. Epigenetic dynamics of monocyte-to-macrophage differentiation. Epigenetics & Chromatin 9, 33, doi:10.1186/s13072-016-0079-z (2016).

9 Tirado-Magallanes, R., Rebbani, K., Lim, R., Pradhan, S. & Benoukraf, T. Whole genome DNA methylation: beyond genes silencing. Oncotarget 8, 5629–5637, doi:10.18632/oncotarget.13562 (2017).

10 Schübeler, D. Function and information content of DNA methylation. Nature 517, 321–326, doi:10.1038/nature14192 (2015).

11 Chen, Y. et al. Genome-wide DNA methylation profiling shows a distinct epigenetic signature associated with lung macrophages in cystic fibrosis. Clinical epigenetics 10, 152–152, doi:10.1186/s13148-018-0580-2 (2018).

12 Fricker, M. & Gibson, P. G. Macrophage dysfunction in the pathogenesis and treatment of asthma. European Respiratory Journal 50, 1700196, doi:10.1183/13993003.00196-2017 (2017).

13 He, L.-X., Tang, Z.-H., Huang, Q.-S. & Li, W.-H. DNA Methylation: A Potential Biomarker of Chronic Obstructive Pulmonary Disease. Frontiers in cell and developmental biology 8, 585–585, doi:10.3389/fcell.2020.00585 (2020).

14 Armstrong, D. A. et al. DNA Methylation Changes in Regional Lung Macrophages Are Associated with Metabolic Differences. ImmunoHorizons 3, 274, doi:10.4049/immunohorizons.1900042 (2019).

15 Martinez, F. J. et al. Idiopathic pulmonary fibrosis. Nature Reviews Disease Primers 3, 17074, doi:10.1038/nrdp.2017.74 (2017).

16 Chambers, R. C. & Mercer, P. F. Mechanisms of alveolar epithelial injury, repair, and fibrosis. Annals of the American Thoracic Society 12 Suppl 1, S16–S20, doi:10.1513/AnnalsATS.201410-448MG (2015).

17 Allen, R. J. et al. Genome-Wide Association Study of Susceptibility to Idiopathic Pulmonary Fibrosis. Am J Respir Crit Care Med 201, 564–574, doi:10.1164/rccm.201905-1017OC (2020).

18 Molyneaux, P. L. et al. Host–Microbial Interactions in Idiopathic Pulmonary Fibrosis. American Journal of Respiratory and Critical Care Medicine 195, 1640–1650, doi:10.1164/rccm.201607-1408OC (2017).

19 Vancheri, C., Failla, M., Crimi, N. & Raghu, G. Idiopathic pulmonary fibrosis: a disease with similarities and links to cancer biology. European Respiratory Journal 35, 496, doi:10.1183/09031936.00077309 (2010).

20 Allden, S. J. et al. The Transferrin Receptor CD71 Delineates Functionally Distinct Airway Macrophage Subsets during Idiopathic Pulmonary Fibrosis. American Journal of Respiratory and Critical Care Medicine 200, 209–219, doi:10.1164/rccm.201809-1775OC (2019).

21 Ogger, P. P. et al. Itaconate controls the severity of pulmonary fibrosis. Science Immunology 5, eabc1884, doi:10.1126/sciimmunol.abc1884 (2020).

22 Pidsley, R. et al. Critical evaluation of the Illumina MethylationEPIC BeadChip microarray for whole-genome DNA methylation profiling. Genome Biol 17, 208, doi:10.1186/s13059-016-1066-1 (2016).

23 Stunnenberg, H. G. & Hirst, M. The International Human Epigenome Consortium: A Blueprint for Scientific Collaboration and Discovery. Cell 167, 1145–1149, doi:10.1016/j.cell.2016.11.007 (2016).

24 Okazaki, Y. et al. Increased glucose metabolism in Arid5b-/-skeletal muscle is associated with the down-regulation of TBC1 domain family member 1 (TBC1D1). Biological Research 53, 45, doi:10.1186/s40659-020-00313-3 (2020).

25 Du, Y. et al. Lysophosphatidylcholine acyltransferase 1 upregulation and concomitant phospholipid alterations in clear cell renal cell carcinoma. J Exp Clin Cancer Res 36, 66, doi:10.1186/s13046-017-0525-1 (2017).

26 Javierre, B. M. et al. Lineage-Specific Genome Architecture Links Enhancers and Non-coding Disease Variants to Target Gene Promoters. Cell 167, 1369–1384.e1319, doi:10.1016/j.cell.2016.09.037 (2016).

27 Richeldi, L. et al. Relative versus absolute change in forced vital capacity in idiopathic pulmonary fibrosis. Thorax 67, 407, doi:10.1136/thoraxjnl-2011-201184 (2012).

28 Finucane, O. M., Sugrue, J., Rubio-Araiz, A., Guillot-Sestier, M. V. & Lynch, M. A. The NLRP3 inflammasome modulates glycolysis by increasing PFKFB3 in an IL-1β-dependent manner in macrophages. Sci Rep 9, 4034, doi:10.1038/s41598-019-40619-1 (2019).

29 Herazo-Maya, J. D. et al. Validation of a 52-gene risk profile for outcome prediction in patients with idiopathic pulmonary fibrosis: an international, multicentre, cohort study. The Lancet Respiratory Medicine 5, 857–868, doi:https://doi.org/10.1016/S2213-2600(17)30349-1 (2017).

30 Herazo-Maya, J. D. et al. Peripheral Blood Mononuclear Cell Gene Expression Profiles Predict Poor Outcome in Idiopathic Pulmonary Fibrosis. Science Translational Medicine 5, 205ra136, doi:10.1126/scitranslmed.3005964 (2013).

31 Scott, M. K. D. et al. Increased monocyte count as a cellular biomarker for poor outcomes in fibrotic diseases: a retrospective, multicentre cohort study. The Lancet Respiratory Medicine 7, 497–508, doi:10.1016/S2213-2600(18)30508-3 (2019).

32 Titus, A. J., Gallimore, R. M., Salas, L. A. & Christensen, B. C. Cell-type deconvolution from DNA methylation: a review of recent applications. Human molecular genetics 26, R216–R224, doi:10.1093/hmg/ddx275 (2017).

33 Yang, I. V. et al. Relationship of DNA Methylation and Gene Expression in Idiopathic Pulmonary Fibrosis. American Journal of Respiratory and Critical Care Medicine 190, 1263–1272, doi:10.1164/rccm.201408-1452OC (2014).

34 Sanders, Y. Y. et al. Altered DNA Methylation Profile in Idiopathic Pulmonary Fibrosis. American Journal of Respiratory and Critical Care Medicine 186, 525–535, doi:10.1164/rccm.201201-0077OC (2012).

35 Yang, I. V. & Schwartz, D. A. Epigenetics of idiopathic pulmonary fibrosis. Translational research: the journal of laboratory and clinical medicine 165, 48–60, doi:10.1016/j.trsl.2014.03.011 (2015).

36 Bell, C. G. et al. DNA methylation aging clocks: challenges and recommendations. Genome Biology 20, 249, doi:10.1186/s13059-019-1824-y (2019).

37 Horvath, S. & Raj, K. DNA methylation-based biomarkers and the epigenetic clock theory of ageing. Nature Reviews Genetics 19, 371–384, doi:10.1038/s41576-018-0004-3 (2018).

38 Shchukina, I. et al. Epigenetic aging of classical monocytes from healthy individuals. bioRxiv, 2020.2005.2010.087023, doi:10.1101/2020.05.10.087023 (2020).

39 Corces, M. R. et al. Lineage-specific and single-cell chromatin accessibility charts human hematopoiesis and leukemia evolution. Nature Genetics 48, 1193–1203, doi:10.1038/ng.3646 (2016).

40 Okuno, Y., Inoue, K. & Imai, Y. Novel insights into histone modifiers in adipogenesis. Adipocyte 2, 285–288, doi:10.4161/adip.25731 (2013).

41 Claussnitzer, M. et al. FTO Obesity Variant Circuitry and Adipocyte Browning in Humans. New England Journal of Medicine 373, 895–907, doi:10.1056/NEJMoa1502214 (2015).

42 Baba, A. et al. PKA-dependent regulation of the histone lysine demethylase complex PHF2-ARID5B. Nature Cell Biology 13, 668–675, doi:10.1038/ncb2228 (2011).

43 Cichocki, F. et al. ARID5B regulates metabolic programming in human adaptive NK cells. The Journal of experimental medicine 215, 2379–2395, doi:10.1084/jem.20172168 (2018).

44 Liu, Y. et al. Blood monocyte transcriptome and epigenome analyses reveal loci associated with human atherosclerosis. Nature Communications 8, 393, doi:10.1038/s41467-017-00517-4 (2017).

45 Chen, X., Hyatt, B. A., Mucenski, M. L., Mason, R. J. & Shannon, J. M. Identification and characterization of a lysophosphatidylcholine acyltransferase in alveolar type II cells. Proc Natl Acad Sci U S A 103, 11724–11729, doi:10.1073/pnas.0604946103 (2006).

46 Bridges, J. P. et al. LPCAT1 regulates surfactant phospholipid synthesis and is required for transitioning to air breathing in mice. The Journal of Clinical Investigation 120, 1736–1748, doi:10.1172/JCI38061 (2010).

47 Bi, J. et al. Oncogene Amplification in Growth Factor Signaling Pathways Renders Cancers Dependent on Membrane Lipid Remodeling. Cell Metabolism 30, 525–538.e528, doi:https://doi.org/10.1016/j.cmet.2019.06.014 (2019).

48 Xu, Y. et al. Single-cell RNA sequencing identifies diverse roles of epithelial cells in idiopathic pulmonary fibrosis. JCI Insight 1, doi:10.1172/jci.insight.90558 (2017).

49 Leslie, J. et al. c-Rel orchestrates energy-dependent epithelial and macrophage reprogramming in fibrosis. Nature Metabolism, doi:10.1038/s42255-020-00306-2 (2020).

50 Tawakol, A. et al. HIF-1α and PFKFB3 Mediate a Tight Relationship Between Proinflammatory Activation and Anerobic Metabolism in Atherosclerotic Macrophages. Arteriosclerosis, thrombosis, and vascular biology 35, 1463–1471, doi:10.1161/ATVBAHA.115.305551 (2015).

51 Müller, F. et al. RnBeads 2.0: comprehensive analysis of DNA methylation data. Genome Biol 20, 55, doi:10.1186/s13059-019-1664-9 (2019).

52 Zhou, W., Laird, P. W. & Shen, H. Comprehensive characterization, annotation and innovative use of Infinium DNA methylation BeadChip probes. Nucleic Acids Research 45, e22–e22, doi:10.1093/nar/gkw967 (2016).

53 Pidsley, R. et al. A data-driven approach to preprocessing Illumina 450K methylation array data. BMC Genomics 14, 293, doi:10.1186/1471-2164-14-293 (2013).

54 Houseman, E. A. et al. Reference-free deconvolution of DNA methylation data and mediation by cell composition effects. BMC Bioinformatics 17, 259, doi:10.1186/s12859-016-1140-4 (2016).

55 Min, J. L., Hemani, G., Davey Smith, G., Relton, C. & Suderman, M. Meffil: efficient normalization and analysis of very large DNA methylation datasets. Bioinformatics 34, 3983–3989, doi:10.1093/bioinformatics/bty476 (2018).

56 Mansell, G. et al. Guidance for DNA methylation studies: statistical insights from the Illumina EPIC array. BMC Genomics 20, 366, doi:10.1186/s12864-019-5761-7 (2019).

57 Peters, T. J. et al. De novo identification of differentially methylated regions in the human genome. Epigenetics Chromatin 8, 6, doi:10.1186/1756-8935-8-6 (2015).

58 Heinz, S. et al. Simple combinations of lineage-determining transcription factors prime cis-regulatory elements required for macrophage and B cell identities. Mol Cell 38, 576–589, doi:10.1016/j.molcel.2010.05.004 (2010).

59 Breeze, C. E. et al. eFORGE v2.0: updated analysis of cell type-specific signal in epigenomic data. Bioinformatics 35, 4767–4769, doi:10.1093/bioinformatics/btz456 (2019).

60 Pageaud, Y., Plass, C. & Assenov, Y. Enrichment analysis with EpiAnnotator. Bioinformatics 34, 1781–1783, doi:10.1093/bioinformatics/bty007 (2018).

61 Afgan, E. et al. The Galaxy platform for accessible, reproducible and collaborative biomedical analyses: 2016 update. Nucleic Acids Research 44, W3–W10, doi:10.1093/nar/gkw343 (2016).

62 Oki, S. et al. ChIP-Atlas: a data-mining suite powered by full integration of public ChIP-seq data. EMBO reports 19, e46255, doi:10.15252/embr.201846255 (2018).

63 Zhou, G. et al. NetworkAnalyst 3.0: a visual analytics platform for comprehensive gene expression profiling and meta-analysis. Nucleic Acids Res 47, W234–W241, doi:10.1093/nar/gkz240 (2019).

64 Aryee, M. J. et al. Minfi: a flexible and comprehensive Bioconductor package for the analysis of Infinium DNA methylation microarrays. Bioinformatics (Oxford, England) 30, 1363–1369, doi:10.1093/bioinformatics/btu049 (2014).

65 Horvath, S. DNA methylation age of human tissues and cell types. Genome Biology 14, 3156, doi:10.1186/gb-2013-14-10-r115 (2013).

66 Hannum, G. et al. Genome-wide methylation profiles reveal quantitative views of human aging rates. Mol Cell 49, 359–367, doi:10.1016/j.molcel.2012.10.016 (2013).

67 Kulakovskiy, I. V. et al. HOCOMOCO: expansion and enhancement of the collection of transcription factor binding sites models. Nucleic Acids Research 44, D116–D125, doi:10.1093/nar/gkv1249 (2016).

