## Supplementary figures and images for "Epigenetic alterations underlie airway macrophage differentiation and phenotype during lung fibrosis"

### Supplementary Figure 1

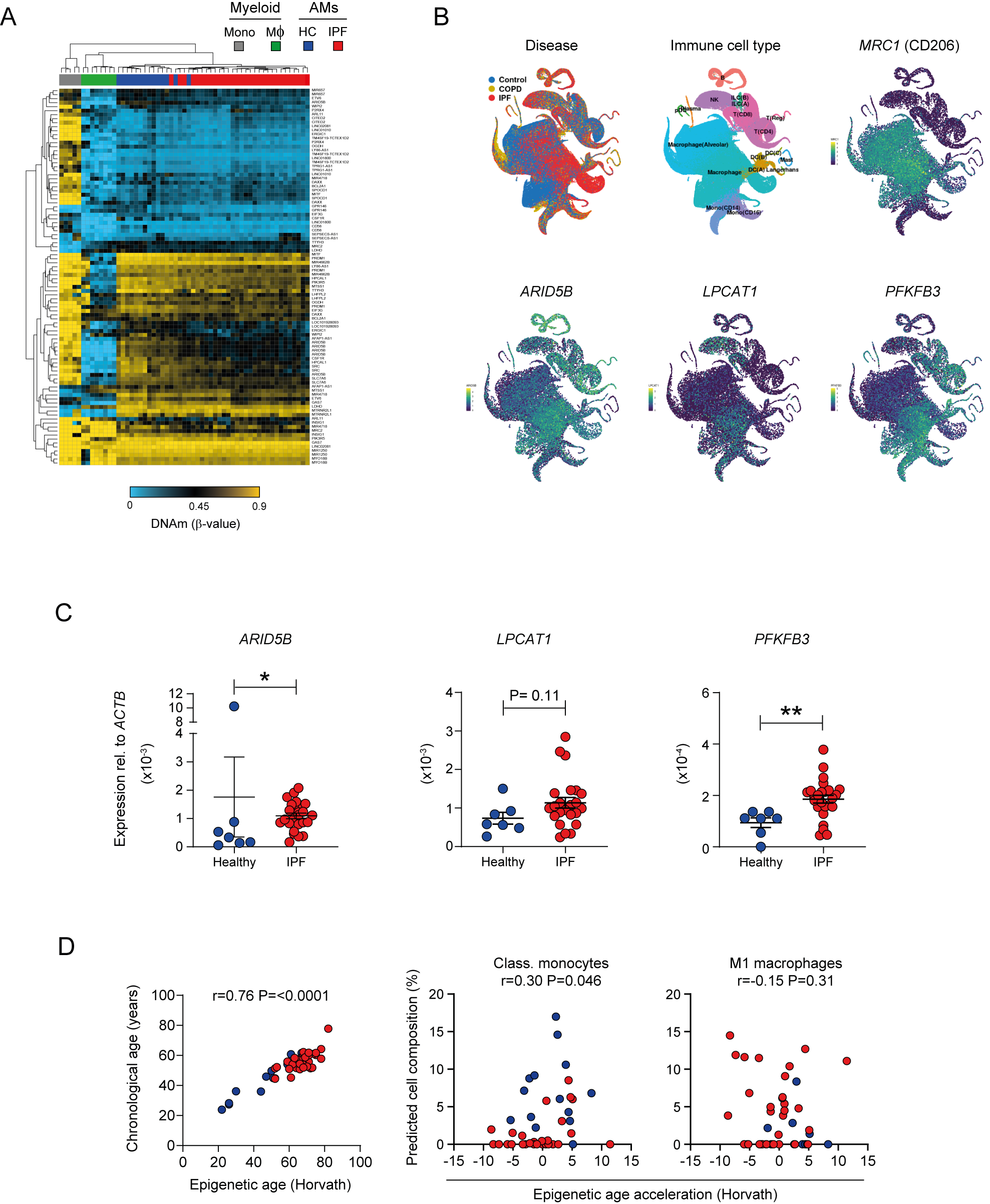

### Supplementary Figure 2

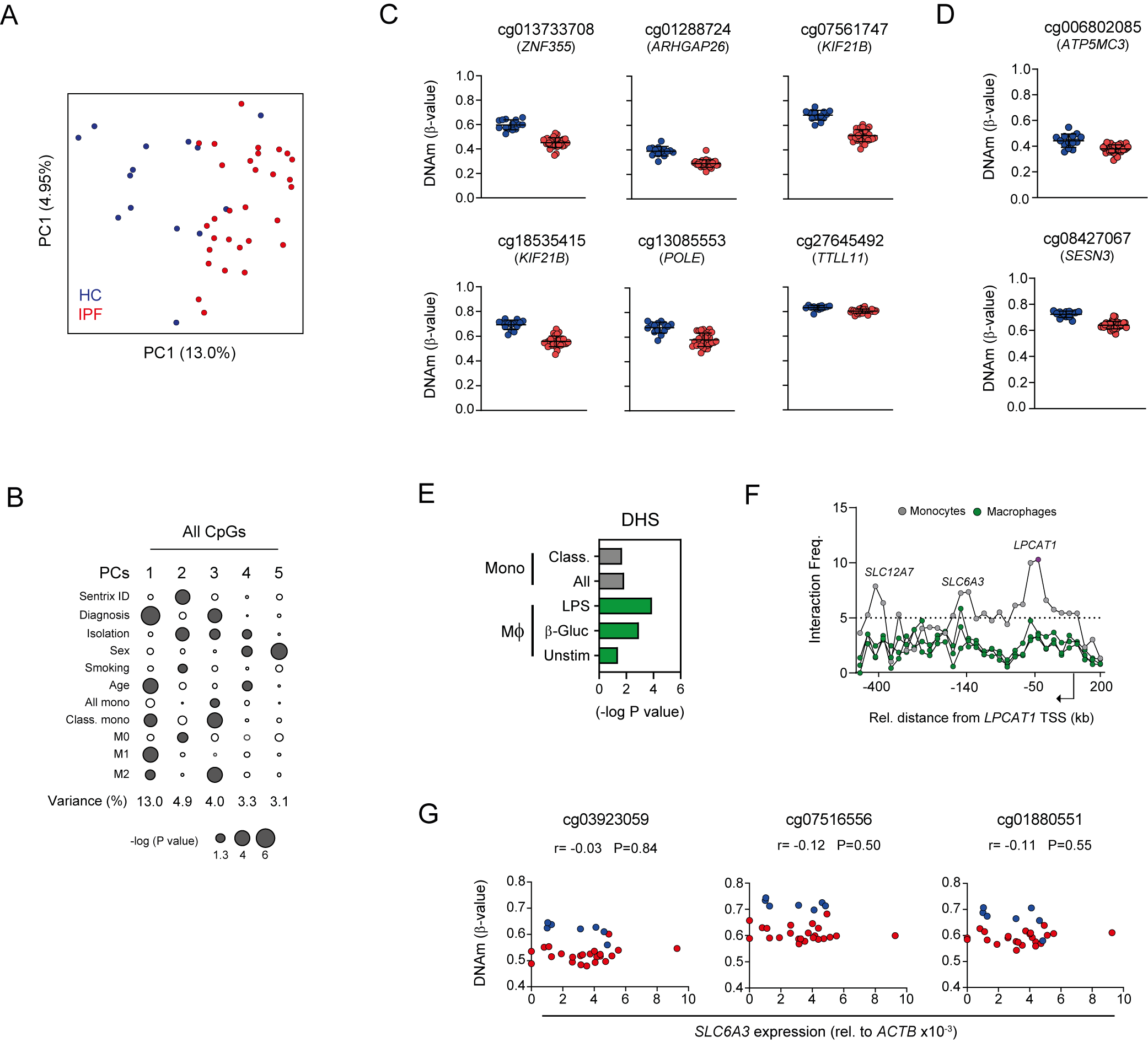

### Supplementary Figure 3

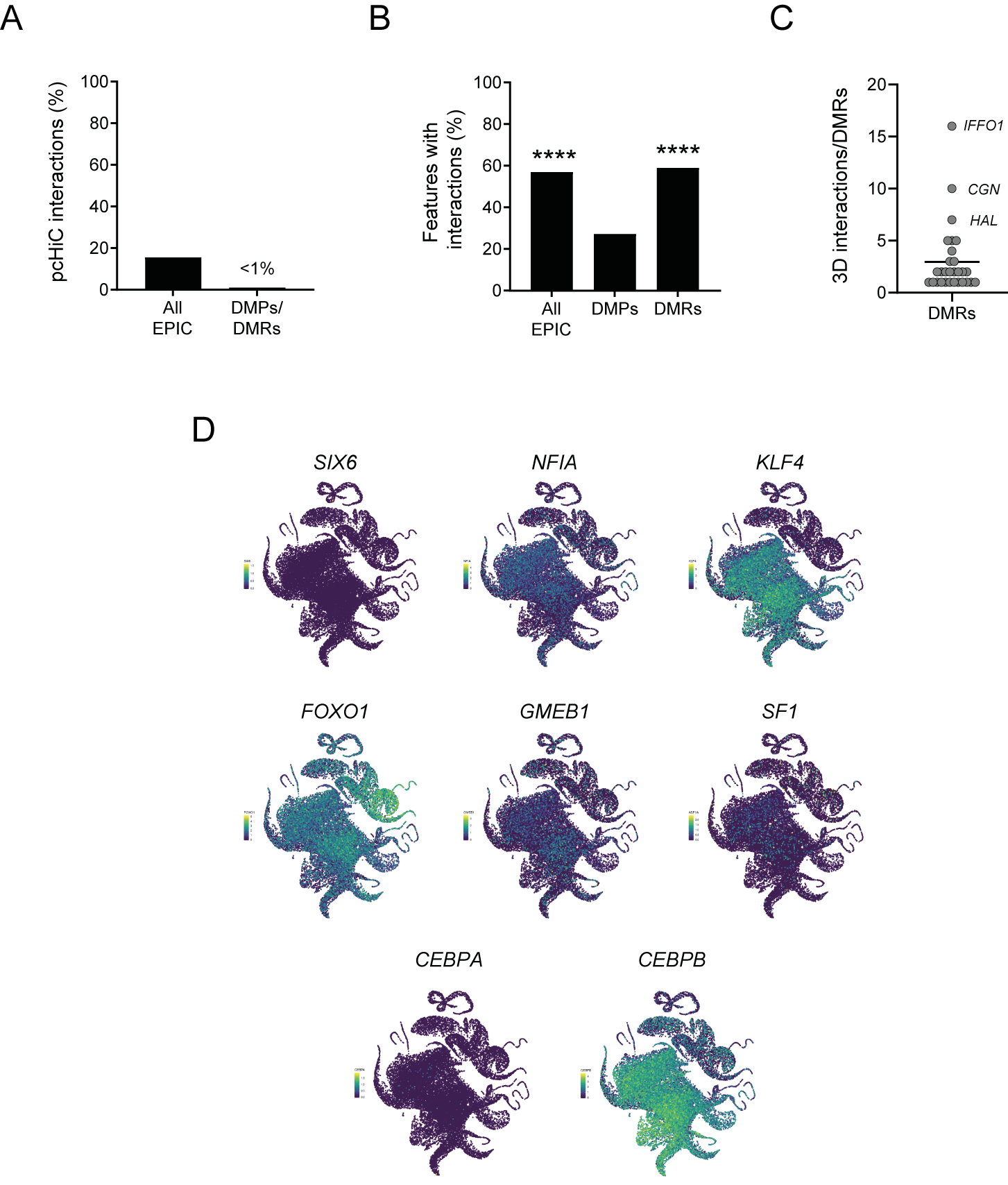

### Supplementary Figure 4

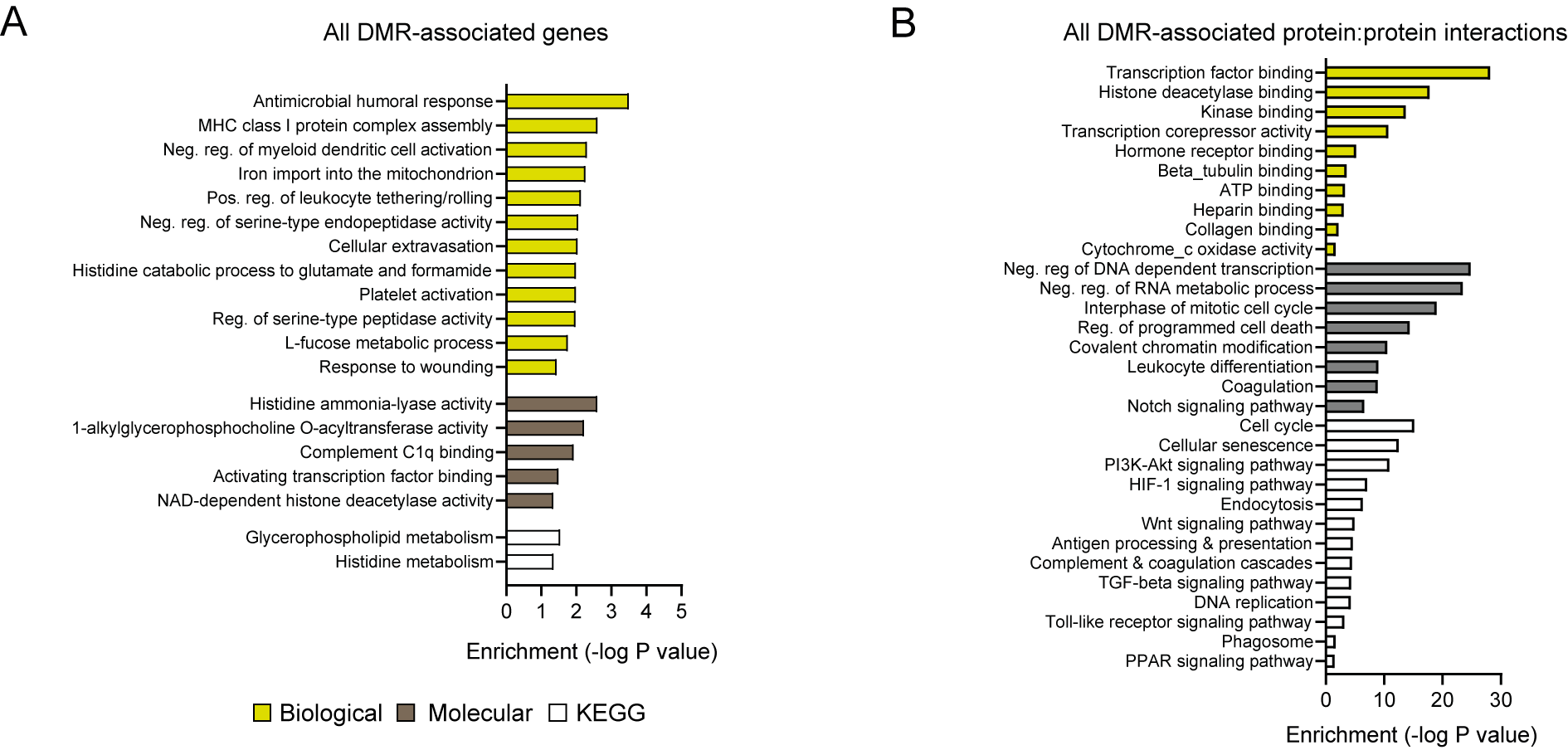
